## Supplementary Figures for "Circulating extracellular vesicle microRNAs mediate immune modulation of social behavior in mice"

### Supplementary Fig. 1

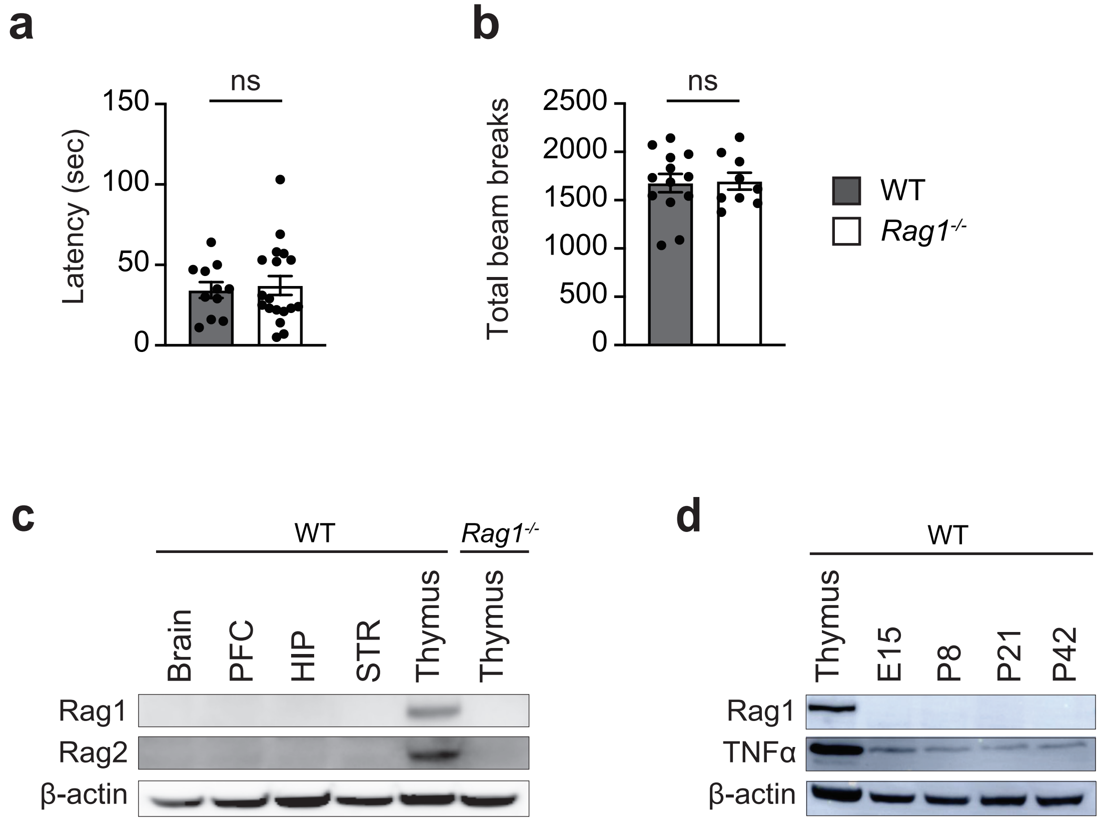

**Supplementary Fig. 1: Additional behavioral data of *Rag1*$^{-/-}$ mice and Rag protein expression in the brain. a**, No difference in latency to find a buried food pellet in the buried food pellet test between WT and *Rag1*$^{-/-}$ mice. WT mice, n = 11 *Rag1*$^{-/-}$ mice, n = 18. **b**, No difference in total activities analyzed with the open field test between WT and *Rag1*$^{-/-}$ mice. WT mice, n = 13 *Rag1*$^{-/-}$ mice, n = 9. **c**, Western blot images showing no detectable Rag1 and Rag 2 proteins in the brains of WT mice. PFC, prefrontal cortex; HC, hippocampus; STR, striatum. **d**, Western blot images showing Rag1, TNFα, and β-actin proteins in the thymus and brains of WT mice (embryonic day 15 (E15), postnatal day 8 (P8), P21, and P42. Rag1 and TNFα images were taken with the same exposure time (5 seconds). Each bar represent mean ± SEM. Each dot represents one mouse. ns, not significant. Significance was determined by Student's *t*-test. See Supplementary Table 13 for the detail of statistical analysis.

### Supplementary Fig. 2

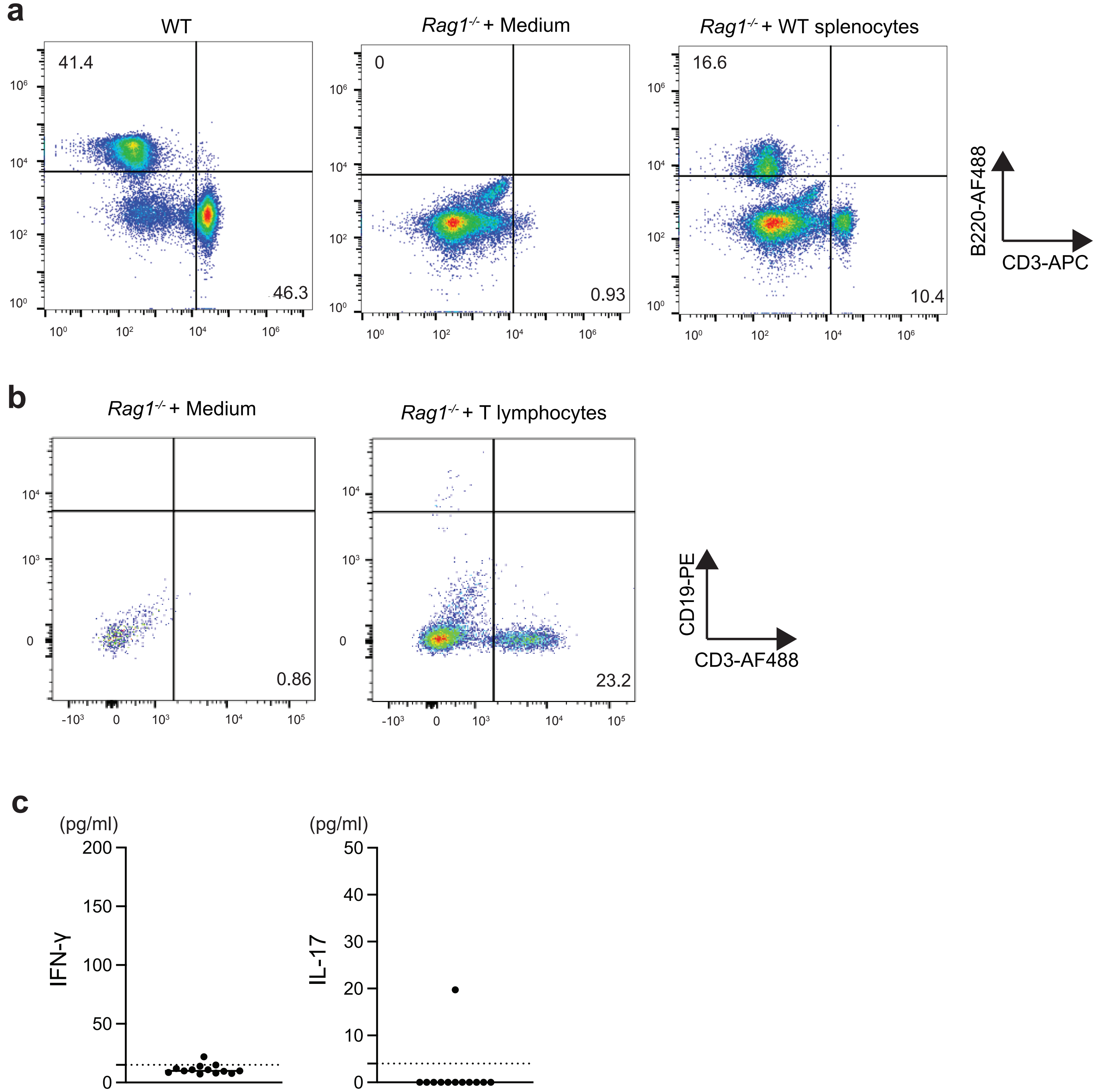

**Supplementary Fig. 2: Lymphocyte reconstitution after adoptive cell transfer**

**a**, Flow cytometry data showing the recovery of $CD3^+$ T cells and $B220^+$ B cells in *Rag1*$^{-/-}$ mice adoptively transferred with WT splenocytes. **b**, Flow cytometry data showing the recovery of $CD3^+$ T cells in *Rag1*$^{-/-}$ mice adoptively transferred with WT T cells. **c**, ELISA data showing the levels of IFN-γ and IL-17 in WT mouse sera.

### Supplementary Fig. 3

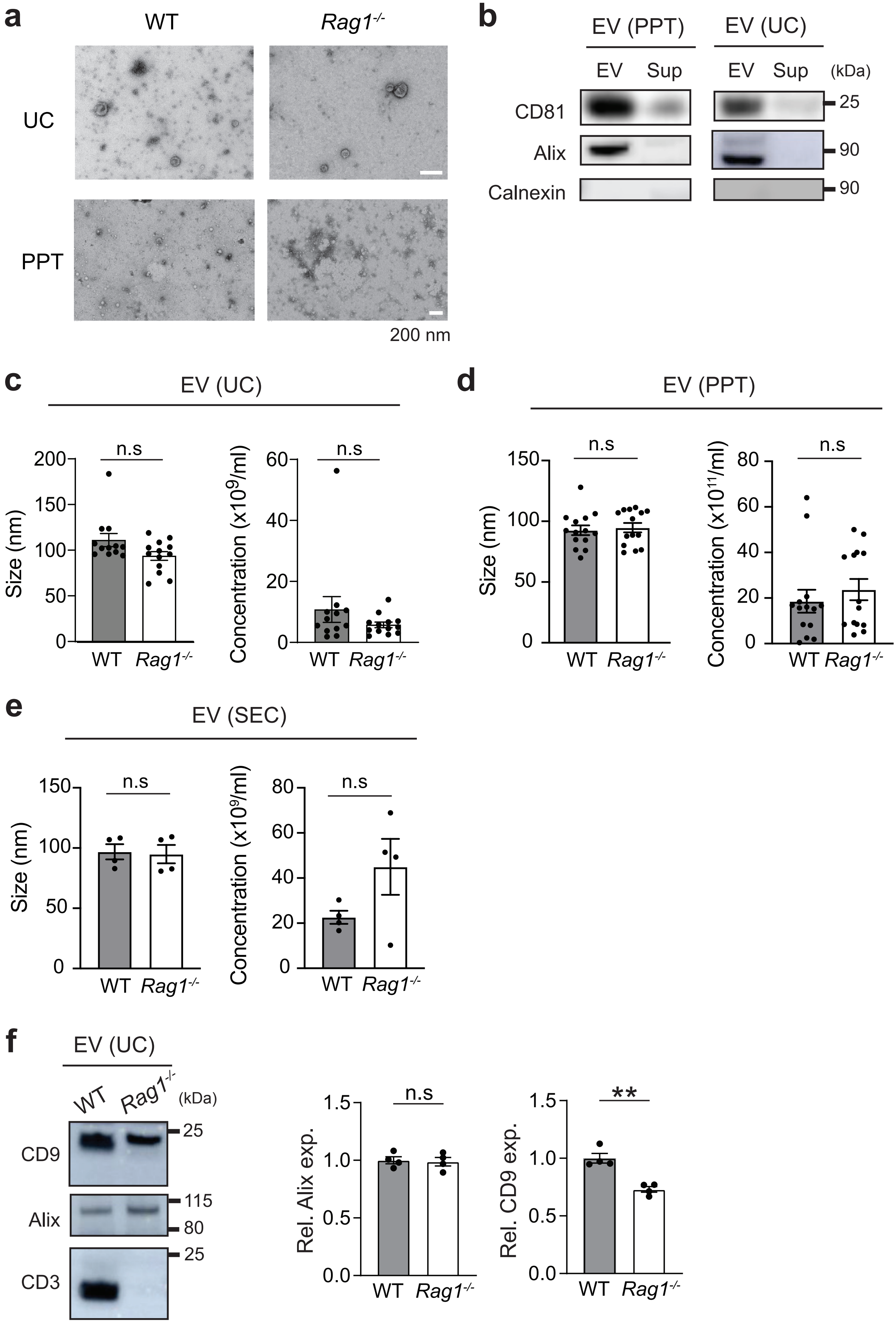

**Supplementary Fig. 3: Characterization of EVs prepared with different methods.**

**a**, Representative transmission electron microscopy (TEM) images of EVs prepared with differential ultracentrifugation (UC) and precipitation (PPT). Scale bar = 200 nm. **b**, Western blot images of EV marker proteins, CD81 and Alix, in EVs prepared with PPT and UC. Calnexin, a cytoplasmic protein marker. Sup, supernatants after EV precipitations. **c**, Nanoparticle tracking analysis of bEVs from WT and $Rag1^{-/-}$ mice using NanoSight NS300. WT,, n = 12; $Rag1^{-/-}$, n = 13. **d**, NanoSight® analysis of bEVs prepared with PPT from WT and $Rag1^{-/-}$ mouse sera. WT, n = 13; $Rag1^{-/-}$, n = 11. **e**, ZetaView® analysis of EVs prepared with size exclusion chromatography (SEC) from WT and $Rag1^{-/-}$ mouse sera. WT, n = 4; $Rag1^{-/-}$, n = 4. **f**, Expression levels of EV marker proteins, CD9 and Alix, in WT and $Rag1^{-/-}$ samples (n=3 per group). Left, representative Western blot images. Right, quantification graphs showing no difference in expression levels of EV marker proteins (CD9 and Alix) between WT and $Rag1^{-/-}$ samples. Equal amounts of proteins were loaded into each lane. Each bar represents mean ± SEM. Each dot represents one mouse. *$p$<0.05, **$p$<0.01. n.s., not significant. Significance was determined by Student's *t* test. See Supplementary Table 13 for the detail of statistical analysis.

### Supplementary Fig. 4

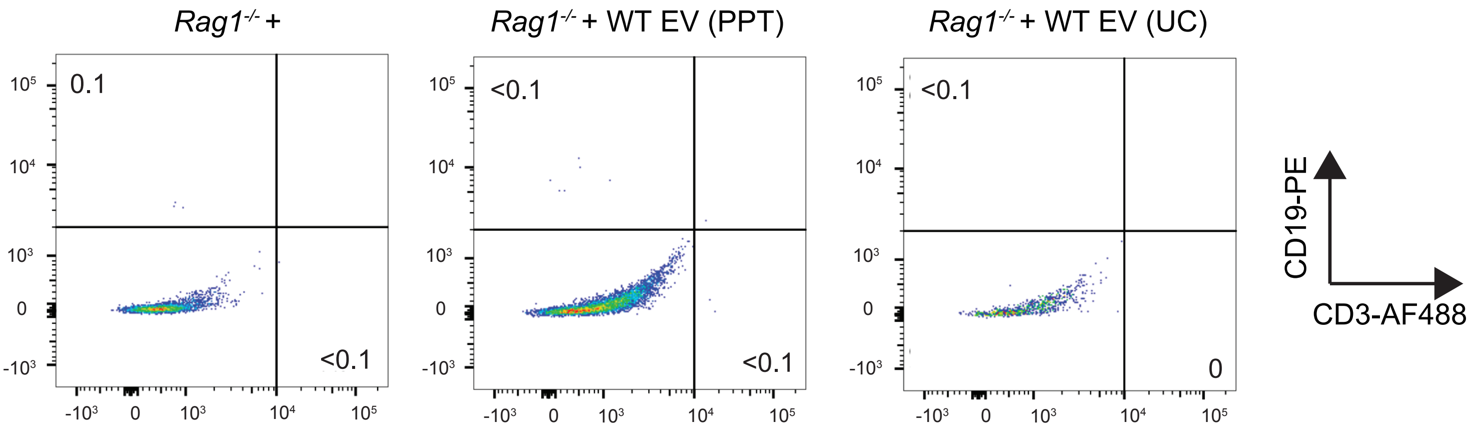

**Supplementary Fig. 4: No recovery of T and B cells in *Rag1*$^{-/-}$ mice intravenously injected with bEVs from WT mice.**

No changes in T or B cell populations were detected in the spleen from *Rag1*$^{-/-}$ mice after intravenous injection with bEVs from WT mice.

### Supplementary Fig. 5

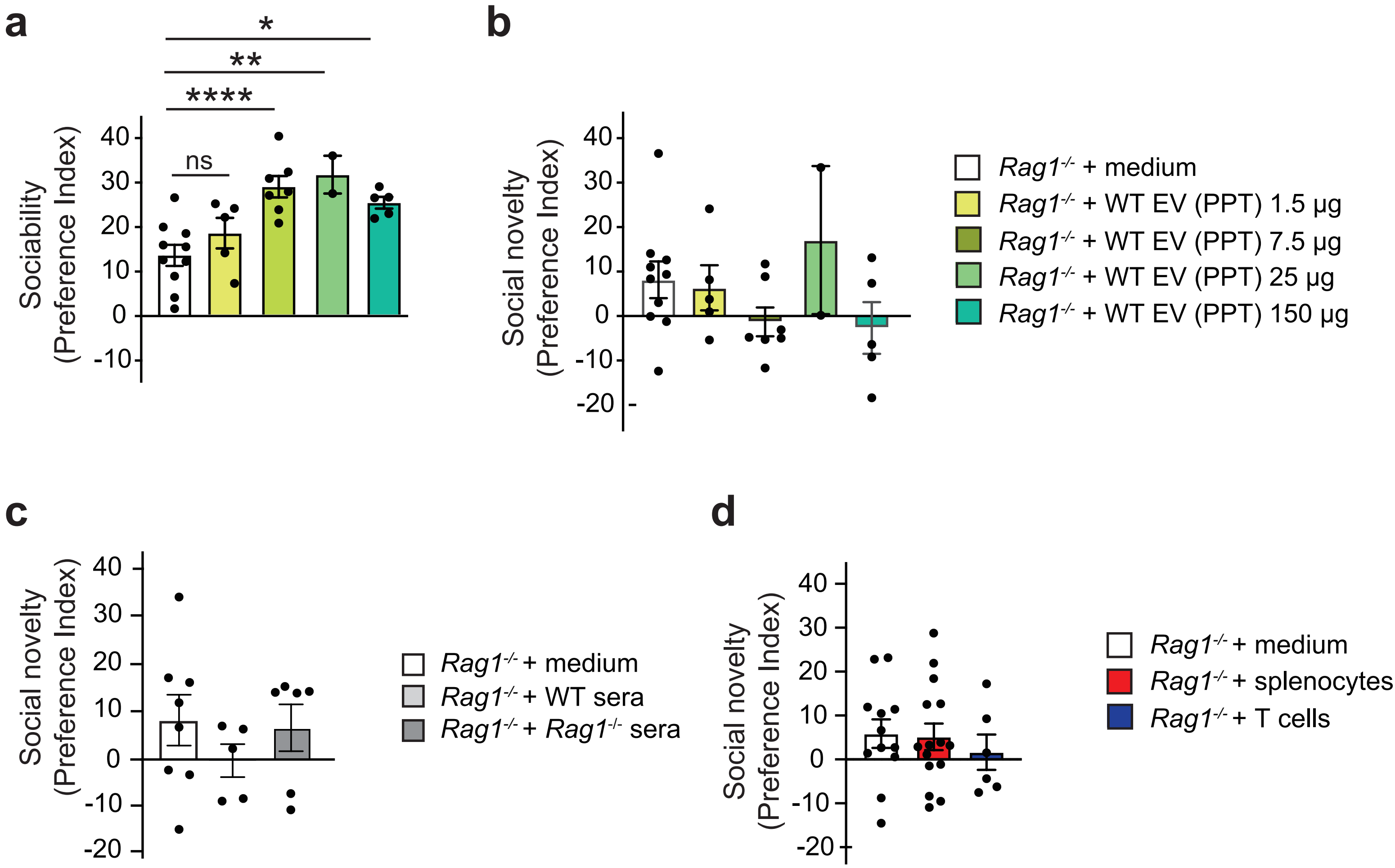

**Supplementary Fig. 5: No recovery in social novelty preference by *Rag1*$^{-/-}$ mice after intravenous injection with WT bEVs or adoptive transfer of WT splenocytes or T cells.**

**a, b**, Intravenous injection of WT bEVs (enriched by PPT) into *Rag1*$^{-/-}$ mice increased their sociability, but not social novelty preference. *Rag1*$^{-/-}$ mice + medium, n = 10; *Rag1*$^{-/-}$ mice + WT bEVs 1.5 μg, n = 5; *Rag1*$^{-/-}$ mice + WT bEVs 7.5 μg, n = 7; *Rag1*$^{-/-}$ mice + WT bEVs 25 μg, n = 2; and *Rag1*$^{-/-}$ mice + WT bEVs 150 μg, n = 5. **c**, Intravenous injection of WT and *Rag1*$^{-/-}$ sera into *Rag1*$^{-/-}$ mice did not show any beneficial effect on their social novelty preference deficits. *Rag1*$^{-/-}$ mice + medium, n = 8; *Rag1*$^{-/-}$ mice + WT sera, n = 5; *Rag1*$^{-/-}$ mice + *Rag1*$^{-/-}$ sera, n = 6. **d**, Adoptive transfer of neither splenocytes nor T cell reconstitution on *Rag1*$^{-/-}$ mice showed any beneficial effect on their social novelty preference deficits. *Rag1*$^{-/-}$ mice + medium, n = 12; *Rag1*$^{-/-}$ mice + WT splenocytes, n = 15; *Rag1*$^{-/-}$ mice + WT T cells, n = 6. Each bar represent mean ± SEM. Each dot represents one mouse. ns, not significant. **p*<0.05, ***p*<0.01, *****p* <0.001. ns, not significant. Significance was determined by one-way ANOVA with *post hoc* Dunnett's test. See Supplementary Table 13 for the detail of statistical analysis.

### Supplementary Fig. 6

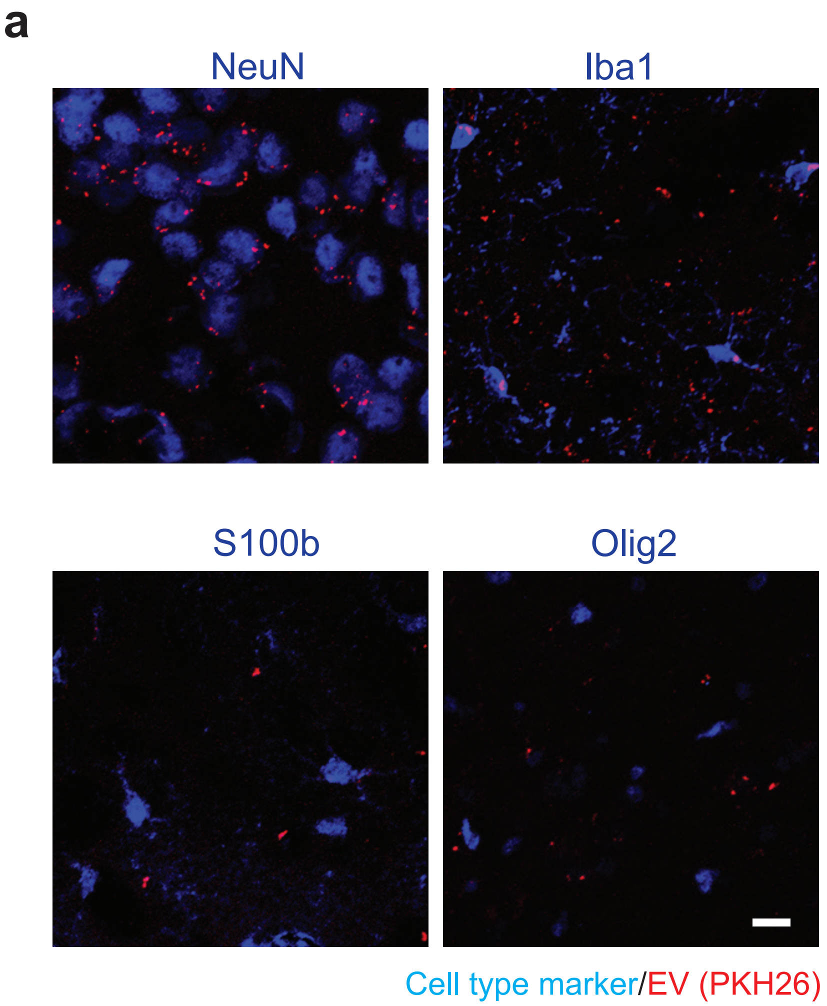


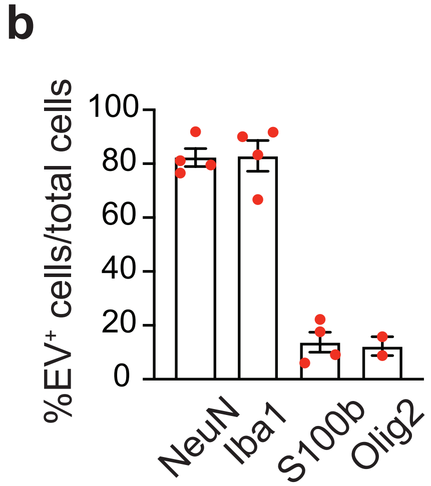

**Supplementary Fig. 6: Co-localization of PKH26-labeled bEVs with neurons and microglia. a,** Representative confocal microscope images showing the localization of PKH26-labeled EVs with $NeuN^+$ neurons and $Iba1^+$ microglia in the mPFC of *Rag1*$^{-/-}$ mice. **b,** Quantification of PKH26-labeled EVs localized with $NeuN^+$ neurons, $Iba1^+$ microglia, $S100b^+$ astrocytes, and $Olig2^+$ oligodendrocyte-lineage cells in the mPFC of *Rag1*$^{-/-}$ mice, n = 3. Scale bar, 10 μm. Each bar represent mean ± SEM.

### Supplementary Fig. 7

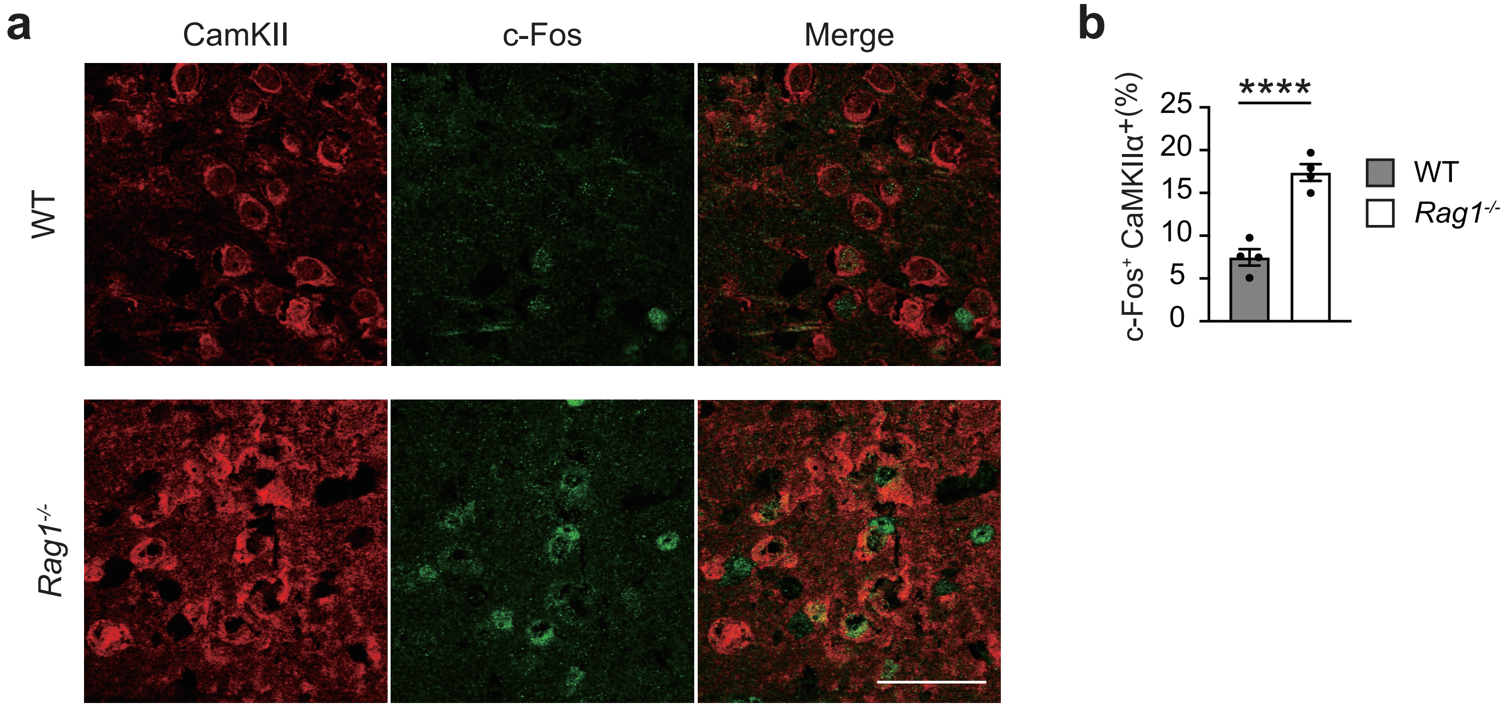

**Supplementary Fig. 7: Increased cFos expression in CaMKIIα$^+$ neurons in the mPFC of *Rag1*$^{-/-}$ mice. a,** Representative images of c-Fos immunoreactivities in CaMKIIα$^+$ neurons in the mPFC of *Rag1*$^{-/-}$ mice compared with WT mice. Scale bar, 50 µm. **b,** Quantification of c-Fos-positive CaMKIIα$^+$ neurons as percentages for c-Fos cells per CaMKIIα$^+$ neurons. WT mice, n = 4 *Rag1*$^{-/-}$ mice, n = 4.

### Supplementary Fig. 8

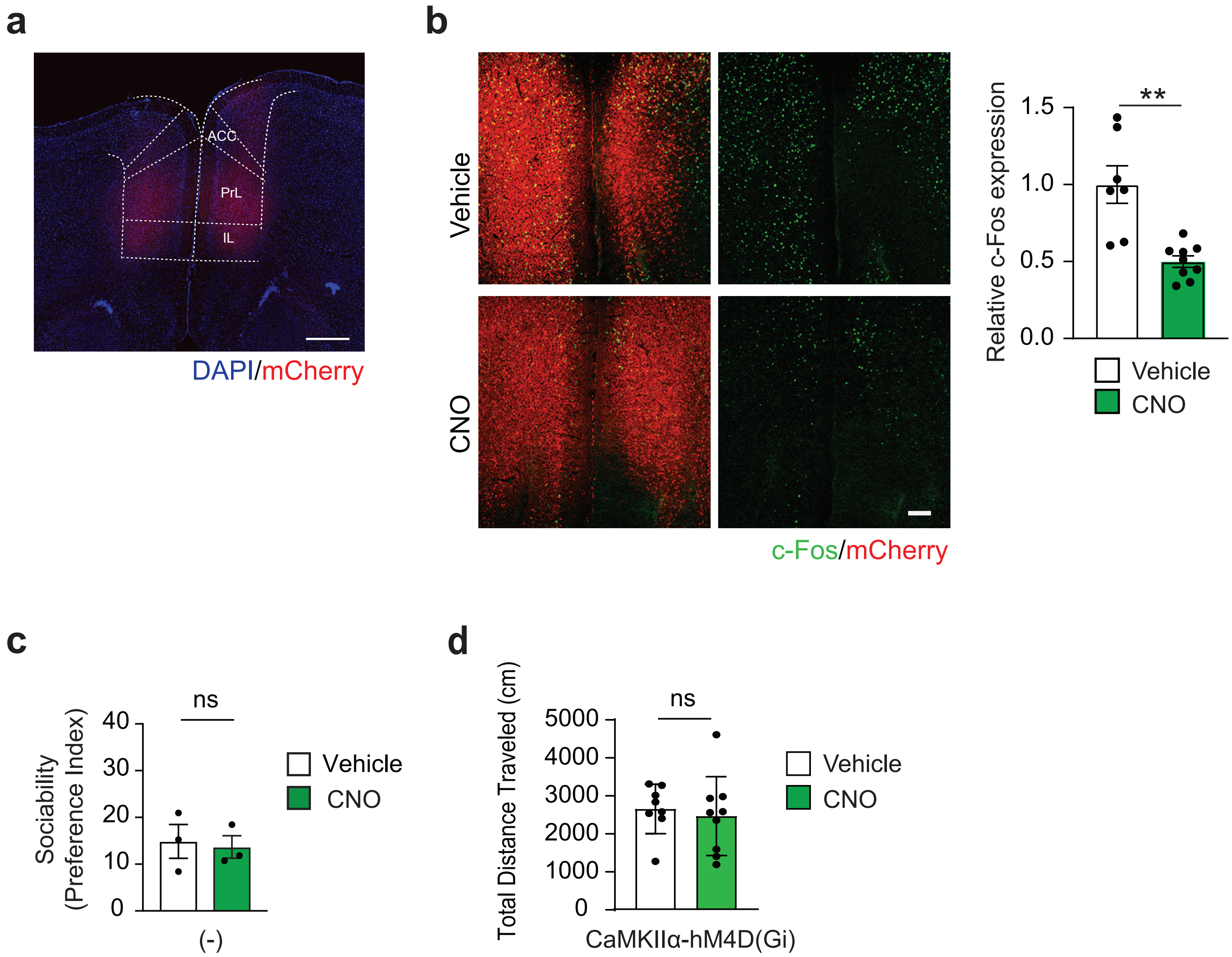

**Supplementary Fig. 8: Validation of DREADD experiments for chemogenic inhibition of CaMKIIα$^{+}$ neurons in the mPFC of *Rag1*$^{-/-}$ mice. a**, Representative image of AAV-CaMKIIα-hM4D(Gi)-mCherry expression in the mPFC of *Rag1*$^{-/-}$ mice. Scale bar, 500 μm. **b**, c-Fos immunoreactivities were suppressed in CNO injected group compared with Vehicle injected group. Vehicle n = 3; CNO n = 3. **c**, No impact of CNO injections on sociability behaviors in *Rag1*$^{-/-}$ mice lacking the expression of DREADD construct. **d**, No significant changes in total travel distance during sociability assays in *Rag1*$^{-/-}$ mice expressing DREADD construct upon CNO injections. Scale bar, 50 μm. Each bar represent mean ± SEM. Each dot represents one mouse. $^{**}p<0.01$. ns, not significant. Significance was determined by Student's *t*-test. See Supplementary Table 13 for the detail of statistical analysis.

### Supplementary Fig. 9

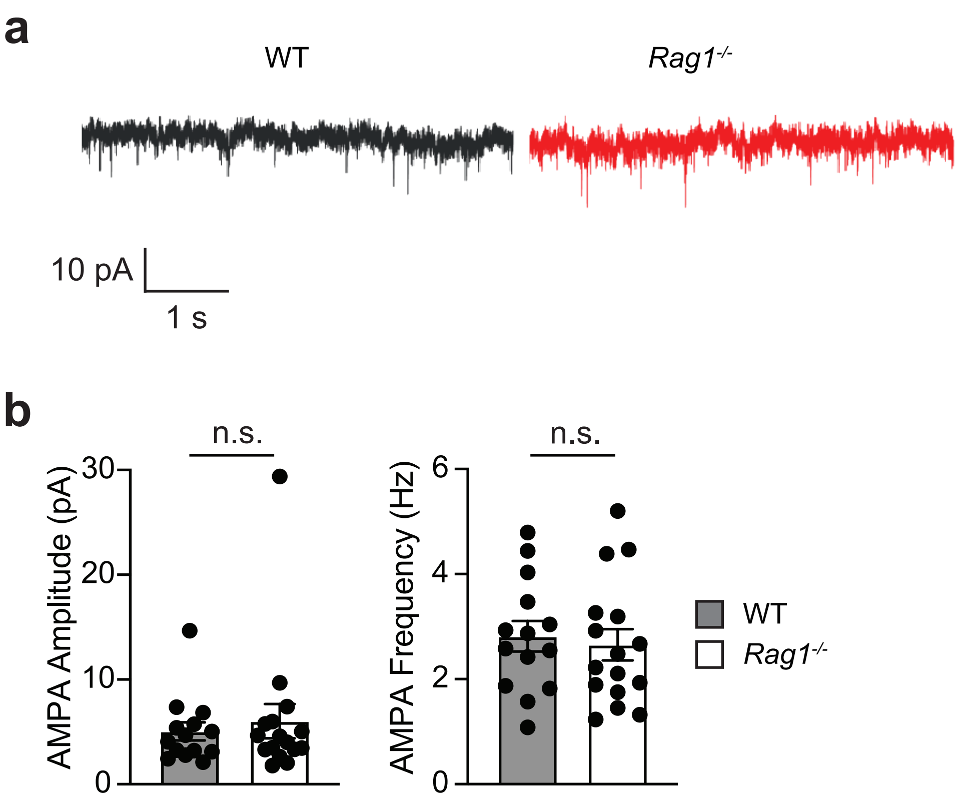

**Supplementary Fig. 9: No changes in spontaneous excitatory AMPAR signaling in the mPFC pyramidal neurons of *Rag1*$^{-/-}$ mice. a,** Representative traces of a gap free recording of pyramidal neuros in the PrL. Recordings were made in the presence of AP-V (50 μM). sEPSCs through AMPA receptors can be seen. **b,** No significant difference in the amplitudes or the frequencies of AMPA currents between WT and *Rag1*$^{-/-}$ mice. Data are mean ± SEM. Sample size was 3 adult male animals per group, with recorded cells used as the statistical unit for analysis. n.s., not significant. Significance was determined by Student's *t*-test. Supplementary Table 13 for the detail of statistical analysis.

### Supplementary Fig. 10

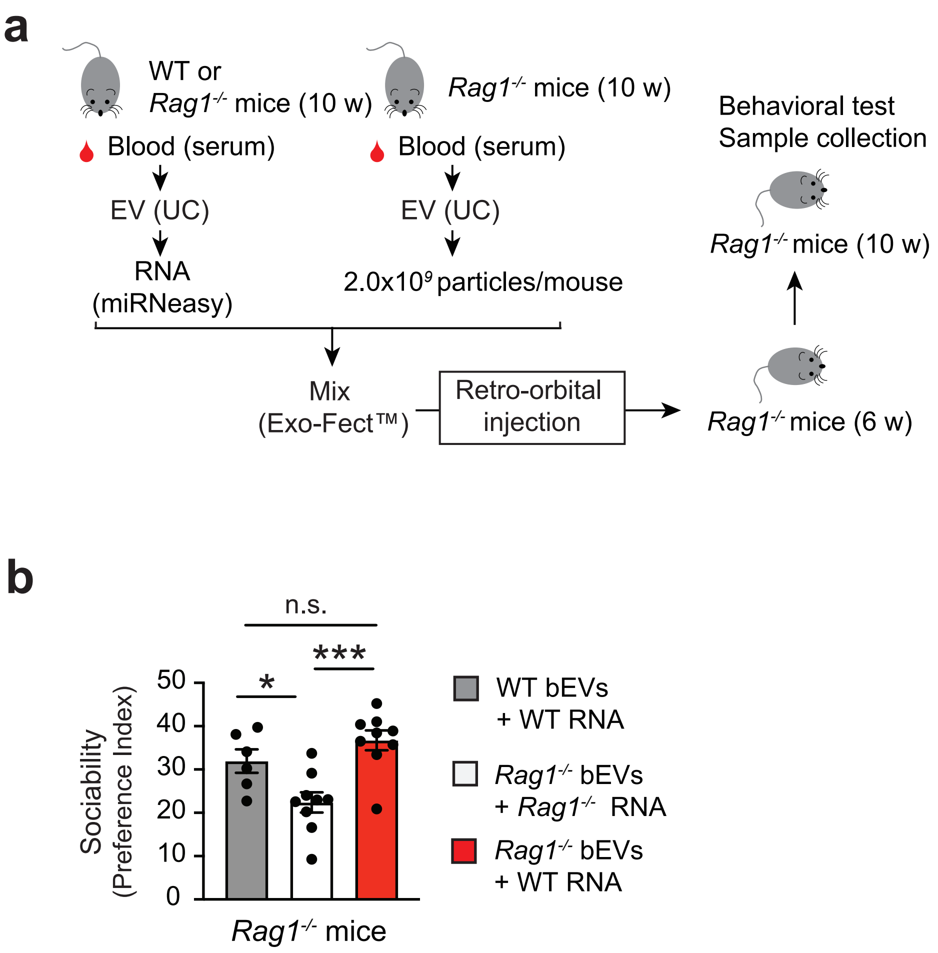

**Supplementary Fig. 10: Differential effects of bEV-derived RNAs.**

**a**, Experimental outline of bEV collection, intravenous injection, and subsequent behavioral assay. **b**, Sociability data of *Rag1*$^{-/-}$ mice administered with WT mice EVs + WT RNA, *Rag1*$^{-/-}$ mice EVs + *Rag1*$^{-/-}$ mice RNA, and *Rag1*$^{-/-}$ mice EVs + WT RNA. WT mice EVs + WT RNA, n = 6; *Rag1*$^{-/-}$ mice EVs + *Rag1*$^{-/-}$ mice RNA, n = 9; *Rag1*$^{-/-}$ mice EVs + WT RNA, n = 9. Each bar represent mean ± SEM. Each dot represents one mouse. *$p$<0.05, ***$p$<0.001. n.s., not significant. Significance was determined by Student's *t*-test. See Supplementary Table 13 for the detail of statistical analysis.

### Supplementary Fig. 11

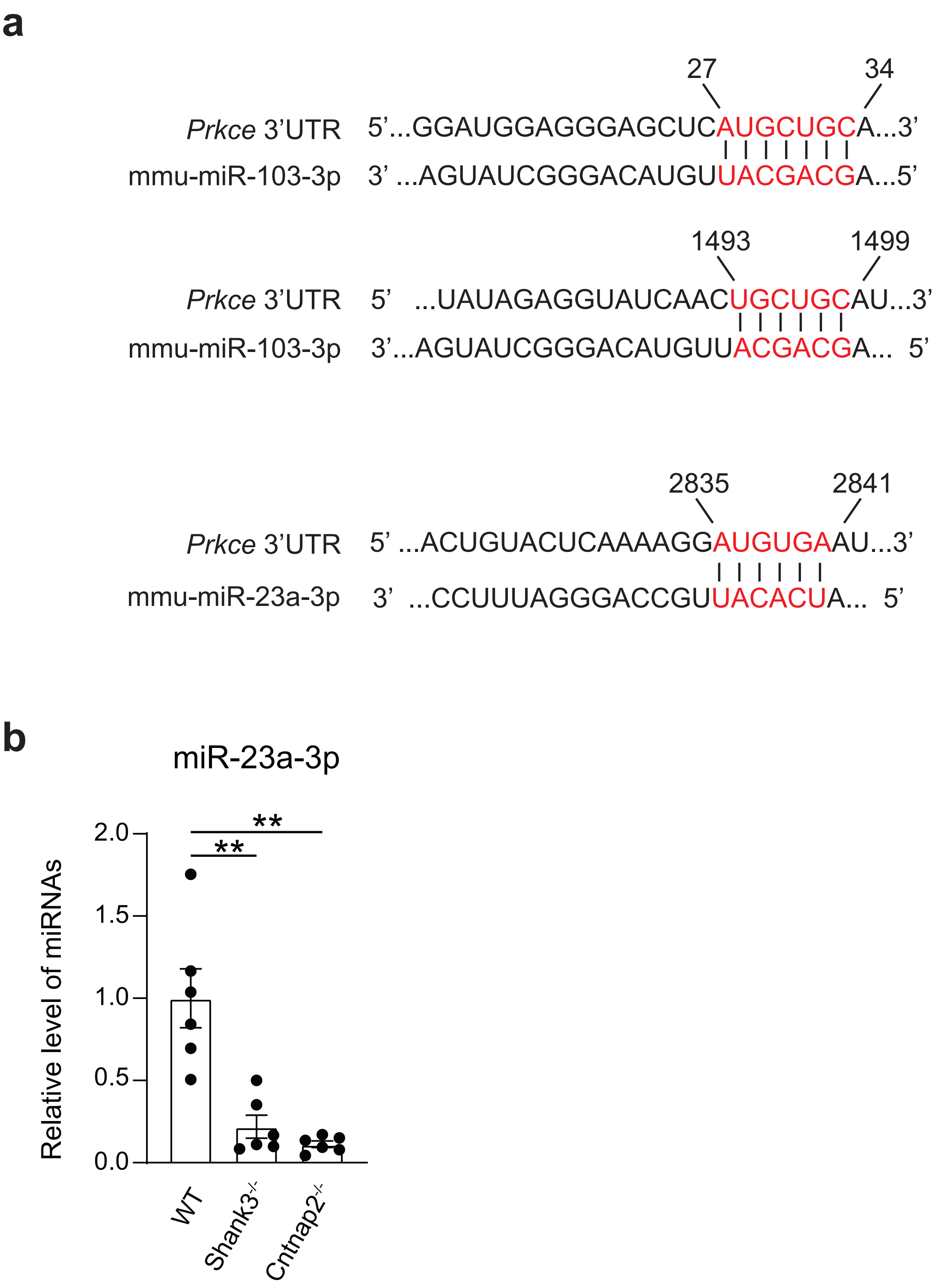

**Supplementary Fig. 11: Predicted targeting sites for miR-23a-3p and miR-103-3p in the 3' UTR of *Prkce* gene.** **a**, Predicted consequential pairings of target region (top) and miRNA (bottom) are shown for miR-23a-3p and miR-103-3p. The target regions are conserved between mouse and human. Prediction data were obtained using TargetScan (release 8.0: September 2021). **b**, Expression of miR-23a-3p in the bEVs from WT, $Shank3^{-/-}$, and $Cntnap2^{-/-}$ mice (n = 6 per group). Data are shown as fold-change relative to WT data. Each bar represent mean ± SEM. ** $p < 0.01$. Each dot represents one mouse. Significance was determined by one-way ANOVA with post hoc Dunnett's test. See Supplementary Table 13 for the detail of statistical analysis.

### Supplementary Fig. 12

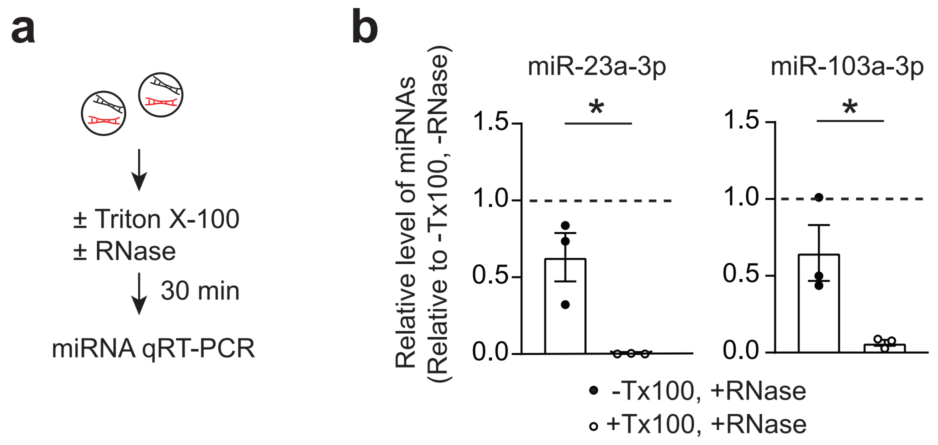

**Supplementary Fig. 12: EVs protect miR-23a-3p and miR-103-3p from RNase.**

**a**, Experimental procedures for Triton-X100 and RNase treatments of bEVs. **b**, Quantitative reverse-transcription PCR (qPCR) data of miR-23a-3p and miR-103-3p expression in WT bEV fractions treated as in **a.** Each bar represent mean ± SEM. $^{**}p<0.01$. Each dot represents one mouse. Significance was determined by Student's *t*-test. See Supplementary Table 13 for the detail of statistical analysis.

### Supplementary Fig. 13

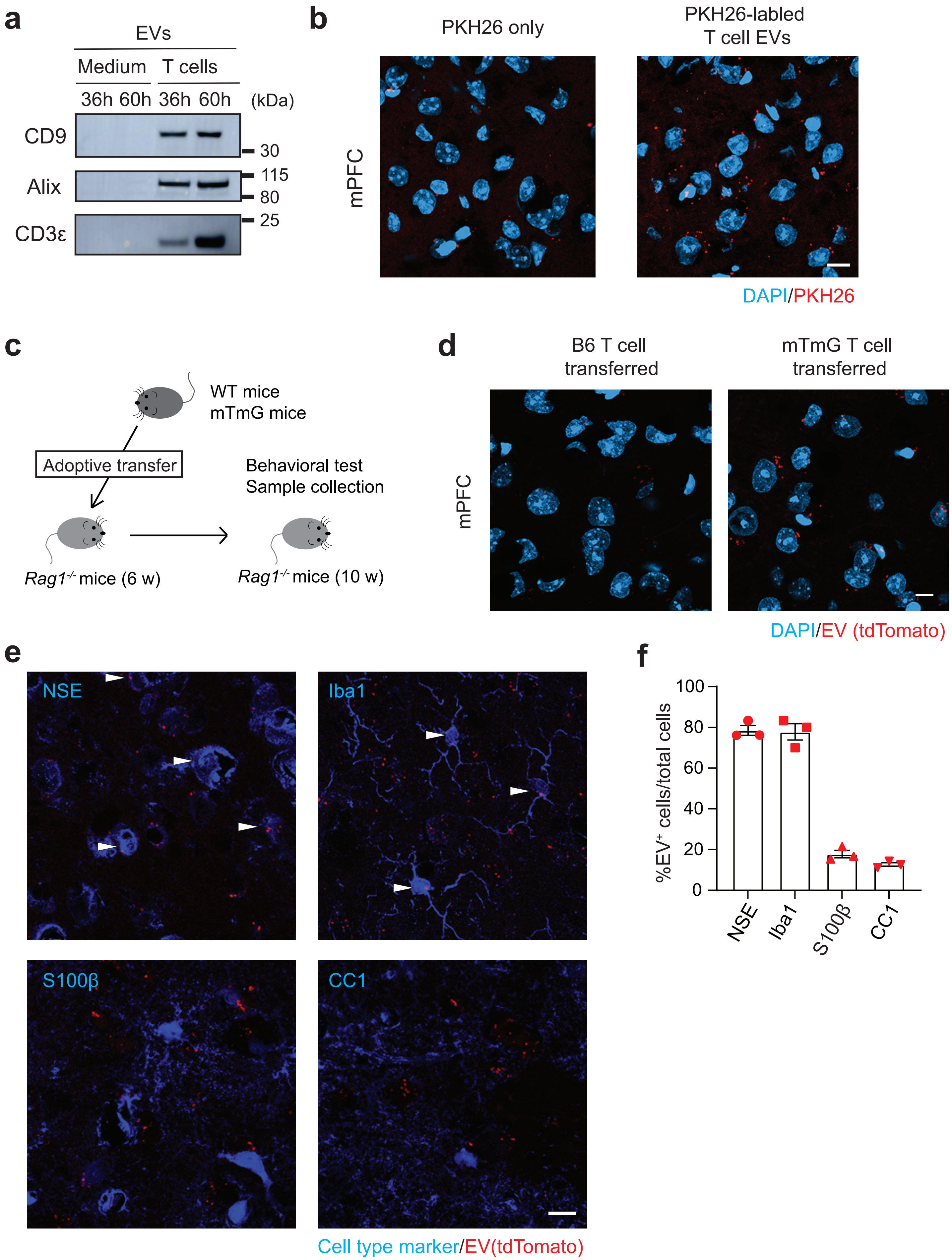

**Supplementary Fig. 13: T cell-derived EVs contain miR-23a-3p and co-localize with neurons following intravenous injection. a**, Western blot data of CD9, Alix, and CD3ε protein expression in EVs harvested from T cell culture medium at 36 h and 60 h after incubation. Culture medium without T cells is used as control. **b**, Representative confocal microscope images of PKH26-labeled T cell-derived EVs (Red) in the mPFC. PKH26 dye injection without EVs are used as control. **c**, Experimental outline of adoptive transfer/serum injection and subsequent behavioral assay. **d**, Representative confocal microscope images of EVs from tdTomato$^+$ T cells adoptively transferred into *Rag1*$^{-/-}$ mice. Immunohistochemistry was conducted 4 weeks after the adoptive transfer. **e**, Representative confocal microscope images of co-localization of tdTomato$^+$ T cell-derived EVs (Red) and neurons, microglia, astrocytes, and oligodendrocytes in the mPFC of *Rag1*$^{-/-}$ mice. EV$^+$ cells are indicated by white arrowheads. **f,** Quantification data of tdTomato$^+$ EV co-localization with specific brain cell types (n = 3 mice). Scale bar, 10 μm. Each bar represent mean ± SEM. Each dot represents one mouse. See Supplementary Table 13 for the detail of statistical analysis.

### Supplementary Fig. 14

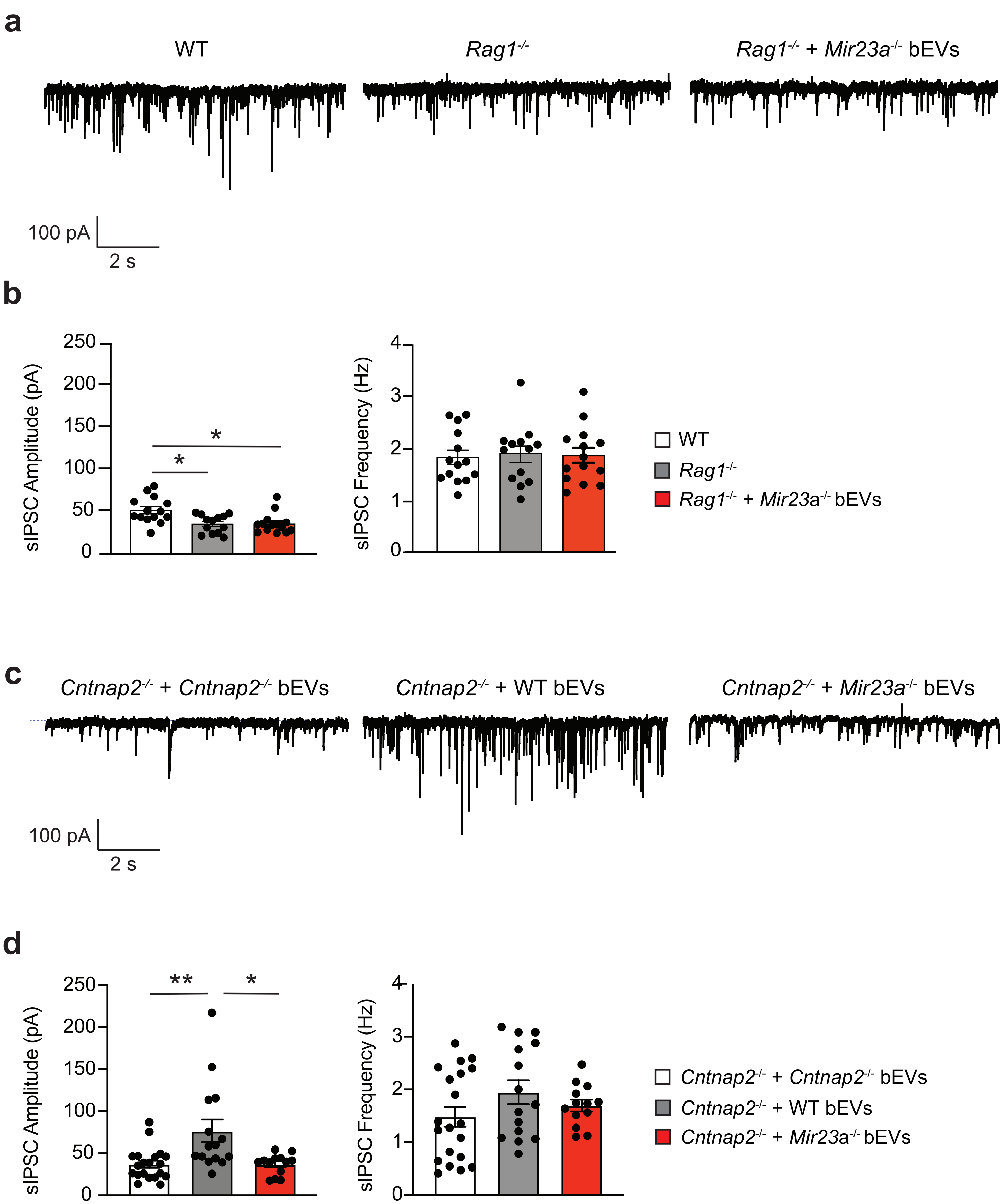

**Supplementary Fig. 14: *Mir23a* deficiency impairs the beneficial effects of bEVs on sIPSCs in mPFC pyramidal neurons of *Rag1*$^{-/-}$ and *Cntnap2*$^{-/-}$ mice. a,** Representative traces of sIPSCs in *Rag1*$^{-/-}$ mPFC (PrL) pyramidal neurons exposed to *Mir23a*$^{-/-}$ bEVs, in comparison to those in WT and *Rag1*$^{-/-}$ mPFC. **b,** Quantification of sIPSC amplitude and frequency between groups. **c,** Representative traces of sIPSCs in *Cntnap2*$^{-/-}$ mPFC (PrL) pyramidal neurons exposed to *Cntnap2*$^{-/-}$, WT and *Mir23a*$^{-/-}$ bEVs as described in Fig. 5. *Cntnap2*$^{-/-}$ + WT bEV data are the same as Fig. 5c, d. **d,** Quantification of sIPSC amplitude and frequency between groups. Each bar represent mean ± SEM. Each dot represents one cell. *$p<0.05$, **$p<0.01$. Significance was determined by Kruskal-Wallis test with *post hoc* Dunn's test. See Supplementary Table 13 for the detail of statistical analysis.
